# Loading-Induced Bone Formation is Mediated by *Wnt1* Induction in Osteoblast-Lineage Cells

**DOI:** 10.1101/2022.02.28.482178

**Authors:** Lisa Y. Lawson, Nicole Migotsky, Christopher J. Chermside-Scabbo, John T. Shuster, Kyu Sang Joeng, Roberto Civitelli, Brendan Lee, Matthew J. Silva

## Abstract

Mechanical loading on the skeleton stimulates bone formation. Although the exact mechanism underlying this process remains unknown, a growing body of evidence indicates that the Wnt signaling pathway is necessary for the skeletal response to loading. Recently, we showed that Wnts produced by osteoblast lineage cells mediate the osteo-anabolic response to tibial loading in adult mice. Here, we report that *Wnt1* specifically plays a crucial role in mediating the mechano-adaptive response to loading. Independent of loading, short-term loss of *Wnt1* in the Osx-lineage resulted in decreased cortical bone area in the tibias of 5-month old mice. In females, strain-matched loading enhanced periosteal bone formation in Wnt1F/F controls, but not in Wnt1F/F; OsxCreERT2 knockouts. In males, strain-matched loading increased periosteal bone formation in both control and knockout mice; however, the periosteal relative bone formation rate was 65% lower in *Wnt1* knockouts versus controls. Together, these findings show that *Wnt1* supports adult bone homeostasis and mediates the bone anabolic response to mechanical loading.

## Introduction

Mechanical loading on the skeleton stimulates bone formation. Although the mechanisms underlying this process are not fully defined, a growing body of evidence indicates that the Wnt signaling pathway plays a role in loading-induced bone formation. For example, ulnar loading suppresses expression of Wnt antagonist Sclerostin, and downregulation of this antagonist is in fact necessary for the bone anabolic response to loading^1–2^. Furthermore, mice lacking Wnt co-receptor Lrp5 exhibit a diminished bone anabolic response to loading, whereas mice with high-bone mass (HBM) mutations in Lrp5 have enhanced responses to loading^3–5^.

Wnt signaling initiates when one of 19 Wnt ligands (‘Wnts’) binds to Frizzled and Lrp5/6 co-receptors^6^. Recently, we reported that blocking Wnts secretion in the Osx-expressing cells of skeletally mature mice reduced the osteo-anabolic response to loading^7^. OsxCreERT2; Wls^F/F^; mice were tamoxifen-treated to inactivate Wntless – the Wnt-specific transport receptor responsible for shuttling Wnt ligands to the cell surface for exocytosis^8^ – thereby blocking the secretion of all Wnts from osteoblasts and osteocytes. We showed that the periosteal response to tibial loading was 65% lower in Wls knockouts relative to WlsF/F controls, indicating that Wnts produced by osteoblast-lineage cells are important mediators of loading-induced bone formation^7^.

Gene expression studies have aimed to identify the key factors that mediate loading-induced bone formation. In our lab and others, RNASeq analysis of cortical bone samples showed that *Wnt1* and *Wnt7b* were differentially expressed in loaded vs non-loaded bones, suggesting that these Wnt ligands in particular may be necessary for the skeletal response to loading^9–11^. Furthermore, age-related declines in the skeletal response to loading are associated with dampened upregulation of *Wnt1* and *Wnt7b* in the bone, while *in vivo* overexpression of *Wnt1* and *Wnt7b* in the osteo-lineage causes increased bone mass^12–14^. Based on these studies, we hypothesized that loading-induced upregulation of *Wnt1* and *Wnt7b* in osteoblasts is necessary for the bone anabolic response to loading. In this communication, we report that blocking *Wnt1* upregulation in the osteoblast-lineage via *Wnt1* deletion in Osx-expressing cells blunts the bone anabolic response to skeletal loading in mice.

## Methods

### Mice

#### Maintenance and genotyping

All animal studies were approved by the Washington University IACUC. Mice were derived from breeders originally gifted by Drs. Henry Kronenberg (OsxCreERT2^15^), Brendan Lee (Wnt1 floxed^13^), or obtained from Jackson Labs (Wnt7b^C3^ floxed; B6; 129X1-Wnt7b^tm2Amc^/J; stock number 008467). Experimental mice were generated by breeding Wnt1F/F dams to OsxCreERT2; Wnt1F/F sires (for *Wnt1* single conditional mutants), or Wnt1F/F; Wnt7bF/F dams to OsxCreERT2; Wnt1F/F; Wnt7F/F sires (for *Wnt1/7b* double conditional mutants). Mice were weaned at 21 days and housed up to five per cage with *ad libitum* access to normal chow and water. Mice were genotyped by Transnetyx (Transnetyx, USA) using tail snip DNA with probes for wild-type *Wnt1* (*Wnt1-1 WT*), conditional *Wnt1* (*L1L2-Bact-P MD*), wild-type *Wnt7b* (*Wnt7b-2 WT*), conditional Wnt7b (*Wnt7b-2 FL*), and OsxCreERT2 (*Cre*). Mice were euthanized by CO_2_ asphyxiation.

#### Tamoxifen induction

Gene deletion was induced in 20 to 22 week old mice. Tamoxifen was dissolved in corn oil to a final concentration of 10mg/ml, and delivered by oral gavage for 5 consecutive days at a dose of 50mg/kg/day. The first day of dosing was “Day 1” in the experimental timeline. Tamoxifen-treated Wnt1F/F and Wnt1F/F; Wnt7bF/F (cre-negative) mice served as genotype controls for *Wnt1* and *Wnt1/7b* knockout experiments, respectively.

### Knockdown validation

Bone-specific deletion was confirmed by DNA recombination PCR and knockdown efficiency was evaluated by RT-qPCR (see Gene Expression below for RT-qPCR methods). Validation experiments were performed 3 weeks after the first tamoxifen dose, on Day 22. Mice used for validation experiments were not subjected to tibial loading.

### Recombination PCR

In preparation for DNA recombination analysis, skeletal and extra-skeletal tissues were crushed manually with mortar and pestle, then digested overnight in Proteinase K. Genomic DNA was isolated using a commercially available kit from Qiagen (DNeasy). 5’-CTGCCCAGCTGGGTTTCTACTACG-3’ (FOR) and 5’-ACCAGCTGCAGACTCTTGGAATCCG-3’ (REV) were used to amplify wild-type Wnt1 DNA from the intact/conditional allele (800bp). 5’-AGTGAGCTAGTACGGGGTCC-3’ (FOR) and 5’-AGGACCATGAACTGATGGCG-3’ (REV) were used to amplify the modified/recombined Wnt1 locus, which produces a *Wnt1* null allele (368bp). 5’-TGACAGAGGATGGGGAGAAG-3’ (FOR) and 5’-GGTCTTTCCAAGGGTGGTCT-3’ (REV) were used to amplify wild-type Wn7b DNA from the intact locus. 5’-GAGGAAGTCAGGCAGGTGTC-3’ (FOR) and 5’-TATCCCACCGATACGCAAAC-3’ (REV) were used to amplify the modified/mutant Wnt7b allele as described previously^16^. PCR reactions were run separately then combined and resolved by electrophoresis on a 2.5% agarose gel.

### Knockdown efficiency qPCR

Tibias were dissected and processed for knockdown efficiency testing as described under *Gene Expression* (RT-qPCR). PrimeTime assay Mm.PT.58.30187381 (Integrated DNA Technologies, USA) was used to assay *Wnt1* expression in the cortical bone. These primers amplify a region spanning exons 2 and 3, which is deleted in knockout tissues. Deletion of exons 2-3 produces a *Wnt1* null allele; thus, qPCR data reflects the abundance of wild-type *Wnt1* mRNA in the bone. *Wnt7b* was assayed using primers spanning exons 2-3: FOR: 3’-CGGGCAAGAACTCCGAGTAG-5’; REV: 3’-GCGACGAGAAAAGTCGATGC-5’. Exon 3 is deleted in the Wnt7b^C3^ conditional mouse.

### Tibial MicroCT

Sequential in vivo microCT scans of the tibia were used to evaluate the effects of short-term *Wnt1* and *Wnt1/7b* deletion on cortical bone morphometry. Tibias were scanned at the mid-diaphysis 24-48 hours before the start of tamoxifen induction (Day 0), followed by a second scan approximately 3 weeks later on Day 22. Under anesthesia (1-3% isoflurane), the left tibia was pulled straight and a 2.1 mm region centered 5 mm proximal to the distal TFJ was scanned on a vivaCT40 with settings optimized for bone (70 kVp, 115 μA, 300 ms integration, 0.4 sigma, 1 support, 1000 projections, 10.5 μm/voxel) (Scanco Medical, Switzerland). Post-scans were completed on mice in situ after sacrifice. For all scans a 1.05 mm region in the center of the scan was contoured in the Scanco software and evaluated for cortical bone area, total area, cortical thickness, polar moment of inertia, and tissue mineral density.

### Tibial loading

Starting on Day 22 after tamoxifen induction, mice were subjected to an *in vivo* tibial loading regimen to stimulate periosteal lamellar bone formation as previously described^17–18^. Briefly, the right tibias of isoflurane-anesthetized mice were positioned vertically between two fixtures and an ElectroPuls E1000 instrument (Instron, USA) was used to apply cyclic axial compression (60 cycles/day, 4Hz) on the right tibia for 5 consecutive days. Contralateral left tibias served as non-loaded controls.

Strain gage analysis was performed on mice that were sacrificed 3 weeks after tamoxifen induction (Day 22). The right tibia was exposed and cleaned of soft tissue before a single-element strain gage was glued to the antero-medial surface 5mm proximal to the distal tibiofibular junction (TFJ). The tibia was axially loaded on a material testing machine (Dynamite 8841) at a peak-to-peak force ranging from −2N to −8N for 12 cycles at 4Hz. LabView data acquisition software was used to collect data, which was analyzed by linear regression to calculate the force-strain equation for each sex-genotype group.

### Strain-matched and force-matched loading

Force-strain equations for each group were used to define the loading force necessary to engender strain-matched loading at a peak target strain of approximately −3000 με. In males, this required a peak force of −10.5 N for both genotypes. In females, strain-matched loading required −9.2 N and −5.4 N in control and knockout mice, respectively. These forces were found to induce a robust lamellar bone formation response in Wnt1F/F males and females (Fig S3). We loaded an additional set of female *Wnt1* knockout mice to −9.2 N (−5600 με) to provide a force-matched comparison with control females, and determine if increasing strain magnitude induced a greater loading response.

### Dynamic histomorphometry

In preparation for bone formation analysis by dynamic histomorphometry, mice were loaded for 5 days (Days 22-26), then dosed with two fluorochromes to label sites of active mineralization. Calcein green (10 mg/kg; Sigma-Aldrich, St. Louis, MO, USA) and alizarin red (30 mg/kg; Sigma-Aldrich) were given by IP injection on Days 26 and 31, respectively, followed by sacrifice on Day 33 (see timeline, Fig 4A). Bilateral tibias (loaded and non-loaded controls) were de-hydrated through ethanol and embedded in MMA plastic as described previously^19^. Transverse sections from the mid-diaphysis (5mm proximal to the TFJ) were used to capture 20µm z-stack images on a Leica confocal microscope (Leica DMi8) for analysis. Bioquant analysis software was used to calculate periosteal (Ps) and endocortical (Ec) MS/BS, MAR, and BFR/BS^20^. Relative bone formation rate (rBFR/BS) was calculated as [loaded – non-loaded]. Relative (*r*) values were used as a net measure of the effect of loading within each animal.

### Gene Expression

Gene expression was analyzed by in situ hybridization and RT-qPCR 3 weeks after tamoxifen induction, 4-hr after the 5th bout of loading.

#### RNAScope in situ hybridization

Tibias from loaded Wnt1F/F mice were fixed for 24 hours in 10% neutral buffered formalin (NBF), de-calcified in 14% EDTA, and submitted to the Musculoskeletal Histology and Morphometry Core (Washington University in St Louis, USA) for paraffin embedding and sectioning. Bones were sectioned to a thickness of 5µm through the transverse plane approximately 5mm distal to the tibial plateau. Hybridization was performed using the HybEZ II Hybridization System (ACDBio) in conjunction with the RNAScope 2.5HD Assay-BROWN kit. Probes for murine *Wnt1*, *Bmp2*, *Bglap, Axin2, Opg, and Id1* were purchased from ACDBio (Catalog No. 401091, 406661, 441211, 488961, and 322335). Sections were counterstained with Hematoxylin and imaged on a Hamamatsu NanoZoomer slide scanning system at the Alafi Neuroimaging Laboratory (Washington University in St. Louis, USA). Expression was quantified by scoring cells within the ROI (site of peak compressive strain) as negative, low, medium, or high. A two-sided Chi-square test was used for statistical analysis.

#### RT-qPCR

Bilateral tibias were collected and snap-frozen in liquid nitrogen as described previously^7, 12^. Briefly, tibias were exposed and soft tissues were removed from the bone surface using forceps, leaving the periosteum intact. Tibias were cut 2mm below the tibial plateau and 1mm below the tibio-fibrular junction to remove the epiphysis and distal tibia. The remainder, consisting largely of compact cortical bone, was spun at 13,000rpm for 30s to remove bone marrow. Bones were pulverized with a Mikro Dismembrator, lysed in Trizol reagent, and phase separated with chloroform reagent. Total RNA was purified using RNeasy Total RNA kit (Qiagen), and cDNA was prepared from samples with RIN > 6.0 using the iScript cDNA Synthesis Kit (BioRad). Gene expression was analyzed with SYBR-based reagents on a StepOne Plus Machine (Applied Biosystems), and reported as a relative expression value (2^-dCT^) relative to reference gene *Tbp*.

### Statistical Analysis

The main effects of genotype (control vs Wnt1Δ or Wnt1/7bΔ) and loading (non-loaded vs loaded) and their interaction were analyzed by 2-factor ANOVA, and *post hoc* Sidak’s multiple comparisons test was used for pairwise comparisons (Prism 7.0, GraphPad Software, Inc., La Jolla, CA, USA). In this analysis the significance of the interaction (“Int”) term is crucial because it indicates whether the effect of loading differed between genotype groups. One-factor ANOVA was used for outcomes where loading was not a factor (eg knockdown efficiency, rBFR/BS). Significance was defined at *p*<0.05, with trends noted at 0.05<*p*<0.10. Individual data points and the mean ± standard deviation are plotted.

## Results

### Tibial loading induces Wnt1 expression in osteocytes

Tibial loading induces *Wnt1* expression in bone^9, 11–12^. To identify the source of Wnt1 in response to loading, *in situ* hybridization was used to evaluate *Wnt1* RNA expression in the loaded and non-loaded tibias of Wnt1F/F control mice after 5 days of loading (Fig 1A). In loaded tibias *Wnt1* expression was detected in 39.3% (± 14.8%) of the osteocytes at the site of peak compressive strain, while in non-loaded control tibias, *Wnt1* expression was detected in only 3.8% (± 2.8%) of the osteocytes in the same region of interest (*p*<0.0001, chi-square test) (Fig 1B). In contrast, little expression was observed in the bone lining cells of either loaded or non-loaded bones after 5 days of loading (Fig S1).

**Figure 1.**
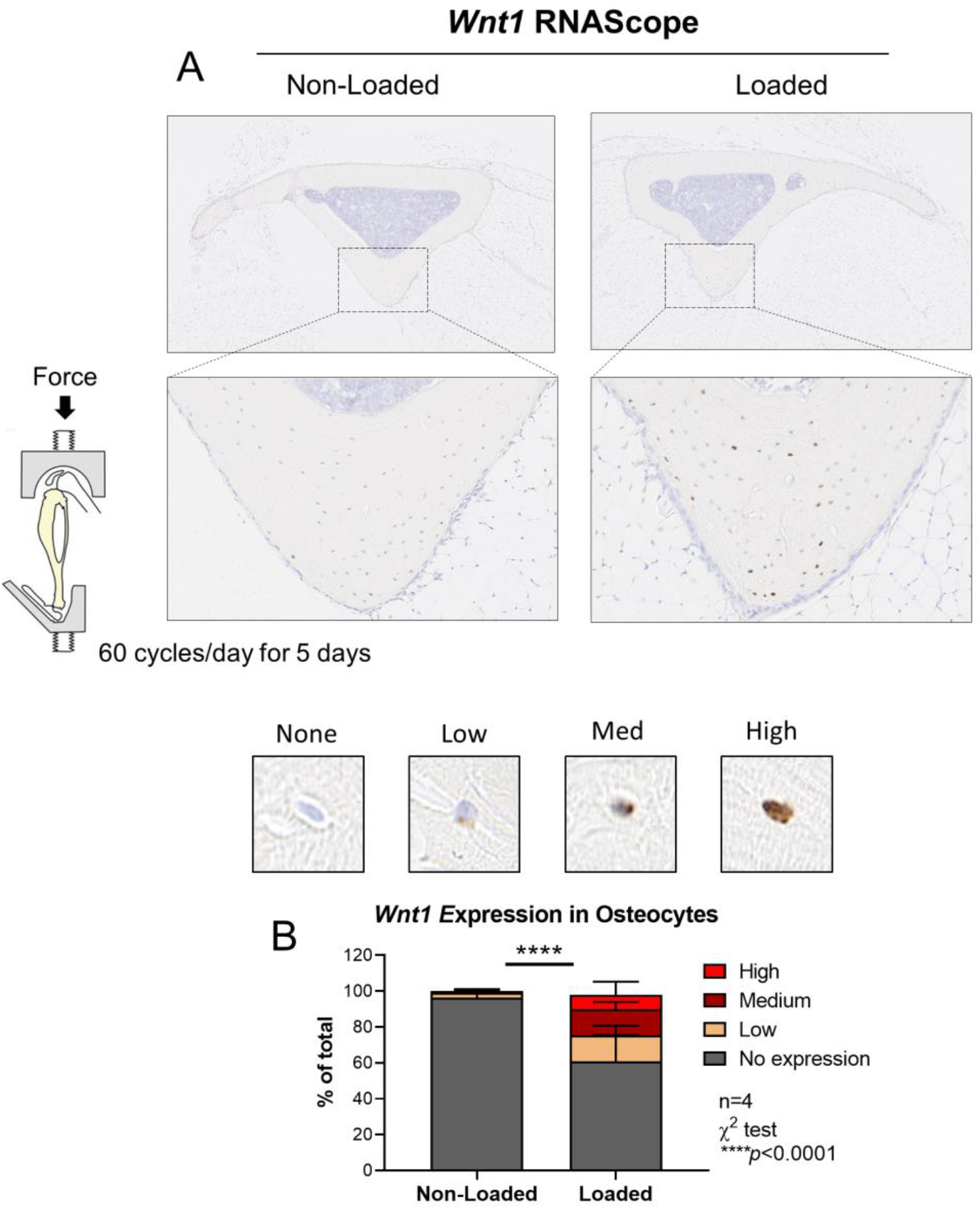
Tibial loading increased *Wnt1* expression in the bone. (A) Tibial *Wnt1* expression was analyzed by RNAScope *in situ* hybridization after 5 days of loading. Results representative of n=4. (B) Osteocytes were scored for *Wnt1* expression at the site of peak compressive strain (lower panels).

### An inducible Cre/LoxP approach was used to delete Wnt1 in the Osx-lineage cells of 5-month ***old mice***

To investigate the role of *Wnt1* in the skeletal response to loading, a tamoxifen-inducible Cre/LoxP approach was used to conditionally inactivate *Wnt1* in the osteoblast-lineage cells of 5-month old mice. OsxCreERT2; Wnt1F/F mice were dosed with tamoxifen by oral gavage for 5 consecutive days to induce exon 2-3 deletion in the osteoblast lineage. Tamoxifen-treated Wnt1F/F mice (Cre-negative) served as genotype controls. Gene deletion was confirmed on Day 22 by DNA recombination PCR and knockdown efficiency testing by RT-qPCR (Fig 2A). DNA recombination PCR indicated that deletion was specific to bone; no deletion occurred in the extra-skeletal tissues surveyed (Fig 2B). PCR analysis also confirmed that bone-specific DNA deletion occurred in 5/5 of the knockout mice tested, but not in any (0/4) of the controls (Fig 2C). Knockdown efficiency testing by qPCR indicated that *Wnt1* mRNA expression was 78% lower in cortical bones of knockouts relative to controls (Fig 2D).

**Figure 2.**
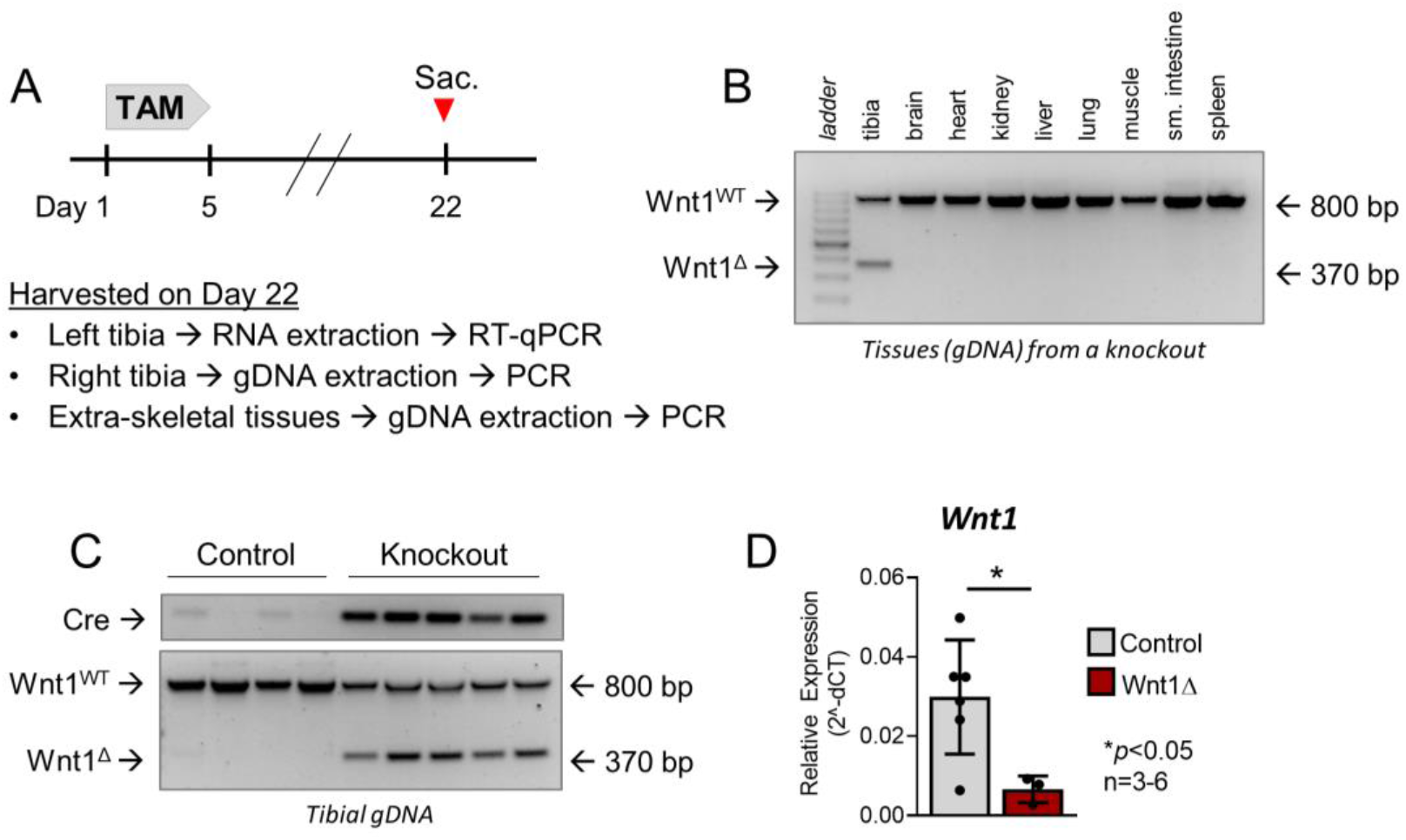
*Wnt1* was conditionally deleted in the Osx-expressing cells of adult mice. (A) Five-month old Wnt1F/F (control) and OsxCreERT2; Wnt1F/F (knockout) mice were dosed with tamoxifen to induce *Wnt1* deletion in the osteoblast lineage. (B-C) DNA Recombination PCR indicated that Wnt1 deletion was specific to the tibias of OsxCreERT2-positive mice. (D) Tibial *Wnt1* RNA expression was 78% lower in *Wnt1* knockouts compared to controls. Individual data points and the mean ± std deviation are shown. n=3-6/group.

### Bone area is decreased after short-term Wnt1 deletion

To evaluate the effects of short-term *Wnt1* deletion on cortical bone morphometry, tibias were serially scanned by *in vivo* microCT 3 weeks apart (Fig 3A). At baseline, there were no differences in cortical morphology between control and *Wnt1* knockout mice, consistent with no phenotype prior to induction of gene deletion. But, 22 days after initial tamoxifen dosing, in both males and females the cortical area and cortical thickness were lower in *Wnt1* knockout mice than in controls, and in females pMOI was also lower in knockouts (Fig. 3B, 3D, 3F). These changes are consistent with a reduction in cortical area in *Wnt1* knockouts from day 0 to 22, whereas cortical area did not change in controls. Similarly, total area was reduced significantly from day 0 to 22 in female knockouts, and marginally in male knockouts (*p*=0.123), whereas it did not change in control mice. In slight contrast, cortical thickness increased from day 0 to 22 in control mice of both sexes, while in knockout mice it was marginally reduced (*p*=0.055, males) or unchanged (*p*>0.05, females). Therefore, *Wnt1* deletion for only 3 weeks led to reduced cortical bone mass, due either to loss of bone or reduced accrual of bone.

**Figure 3.**
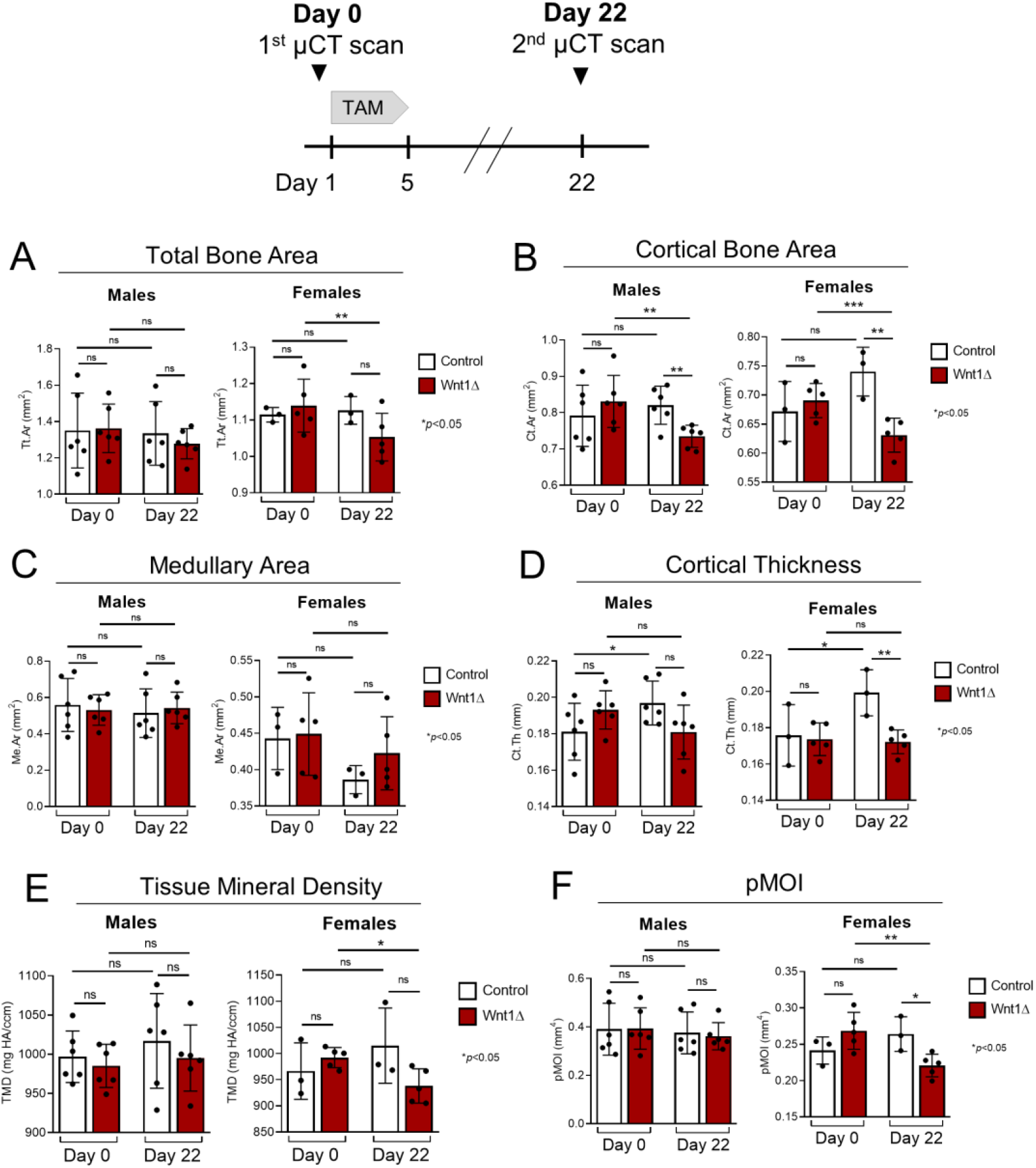
Cortical bone area decreased after short-term *Wnt1* deletion. The tibias of naïve male and female mice were serially scanned to determine the effects of short-term (3 week) *Wnt1* deletion. Tibias were scanned before tamoxifen induction on Day 0, and again on Day 22. Individual data points and the mean ± std deviation are shown. n=3-6/group.

### Loading-induced periosteal bone formation is impaired in Wnt1 knockouts

Next, to evaluate the role of *Wnt1* in the loading response, *in vivo* tibial loading was used to induce bone formation. Three weeks following 5 days of tamoxifen treatment, mice were subjected to daily compressive loading on the right tibia for 5 consecutive days and dosed with calcein and alizarin to label newly mineralizing bone (Fig 4A). Based on *a priori* strain analysis, the majority of mice were subjected to strain-matched loading to a peak target strain of approximately −3000 µε. Preliminary analysis indicated that this stimulus elicited a robust bone formation response in Wnt1F/F controls (Fig S3).

**Figure 4.**
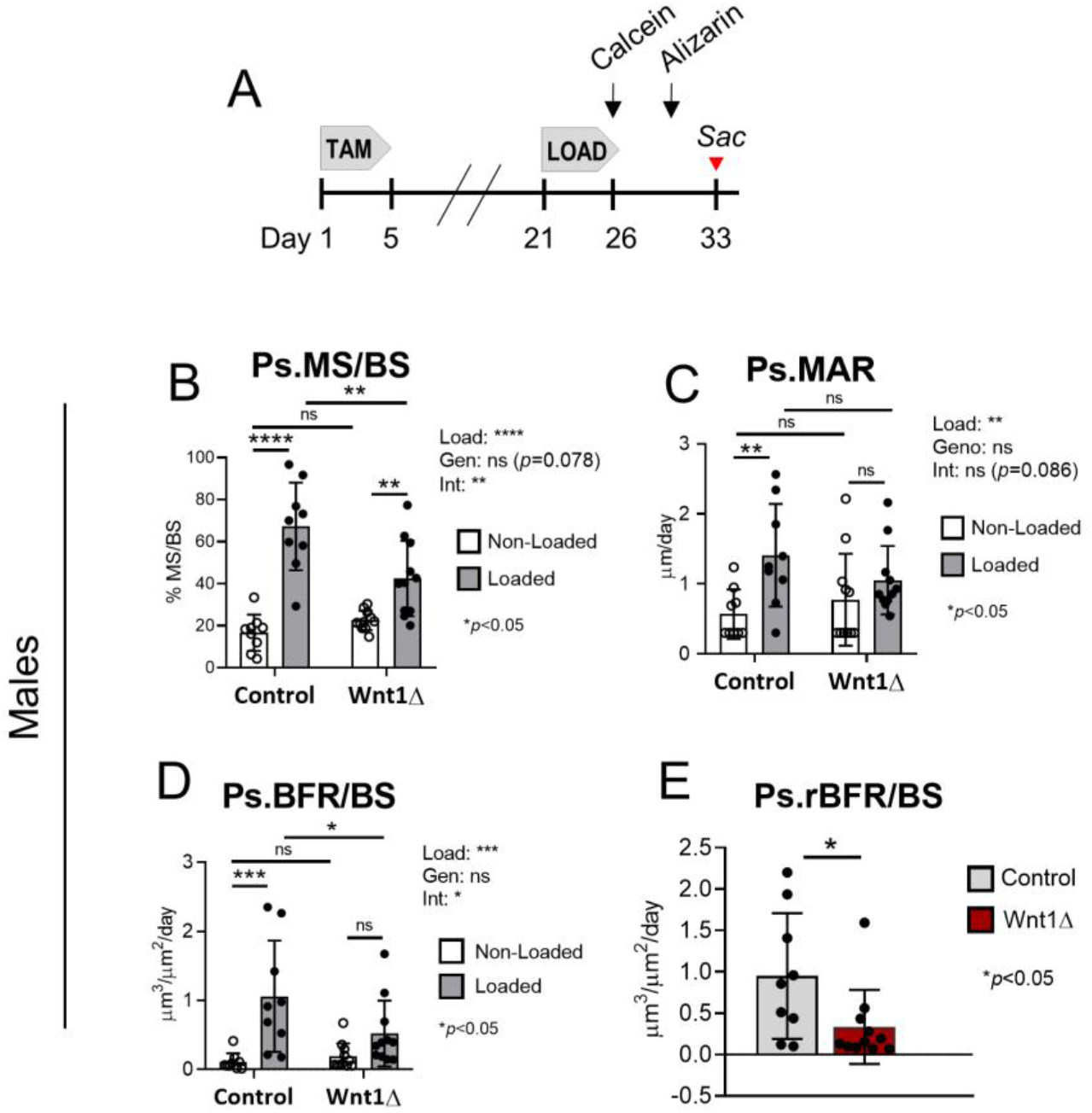
Loading-induced periosteal bone formation was blunted in *Wnt1* knockout males. (A) After 5 days of loading, mice received calcein and alizarin injections in preparation for bone formation analysis by dynamic histomorphometry. (B-D) Periosteal bone formation indices in males subjected to strain-matched loading (−10.5N). Loading increased periosteal bone formation in both groups, albeit to a lesser degree in *Wnt1* knockouts. (E) Relative bone formation rate (rBFR/BS), used as a net index of loading-induced bone formation, was 65% less in *Wnt1* knockouts relative to controls.

In male mice, tibial loading enhanced indices of periosteal (Ps) bone formation in both control and *Wnt1* knockout groups, but the response to loading was diminished in knockouts. Two-way ANOVA indicated a significant effect of loading (main effect) for periosteal bone formation indices, and a significant (or trending) interaction between loading and genotype, indicating that the effect of load was different in control versus *Wnt1* knockout mice (Fig. 4B-D). In controls, Ps.MS/BS, Ps.MAR, and Ps.BFR/BS were 4, 2.5, and 9.6-fold higher, respectively, in loaded vs non-loaded limbs (*p*<0.01) (Fig 4B-4D). By comparison, in knockouts Ps.MS/BS, Ps.MAR, and Ps.BFR/BS were only 1.9 (*p*=0.004), 1.4 (*p*=0.35), and 2.8-fold (*p*=0.16) higher, respectively, in loaded vs. non-loaded limbs. We also calculated the relative bone formation rate (rBFR/BS), defined as BFR/BS_Loaded_ minus BFR/BS_Non-Loaded_, as a measure of loading-induced bone formation. Periosteal rBFR/BS was 65% lower in knockouts relative to controls (*p*=0.037; Fig 4E), which further illustrates that the periosteal response to loading was impaired by the loss of *Wnt1*.

In females, we observed similar negative effects of *Wnt1* deletion on the periosteal response to loading (Fig 5). First, comparing mice loaded to a similar peak strain (∼3000 µƐ; “strain matched”), loading stimulated increases in periosteal bone formation in control but not knockout (Wnt1Δ.) mice. In controls, Ps.MS/BS, Ps.MAR and Ps.BFR/BS were 2.8, 3.5 and 8.3-fold higher, respectively, in the loaded vs non-loaded bones (*p* ≤ 0.0001) (Fig 5A-5C). By comparison, in knockout females loading-associated changes in Ps.MS/BS, Ps.MAR, and Ps.BFR/BS were minimal and not significant. Moreover, periosteal relative bone formation rate (rBFR/BS) was 94% lower in knockouts relative to controls after strain-matched loading (p<0.0001; Fig 5D).

**Figure 5.**
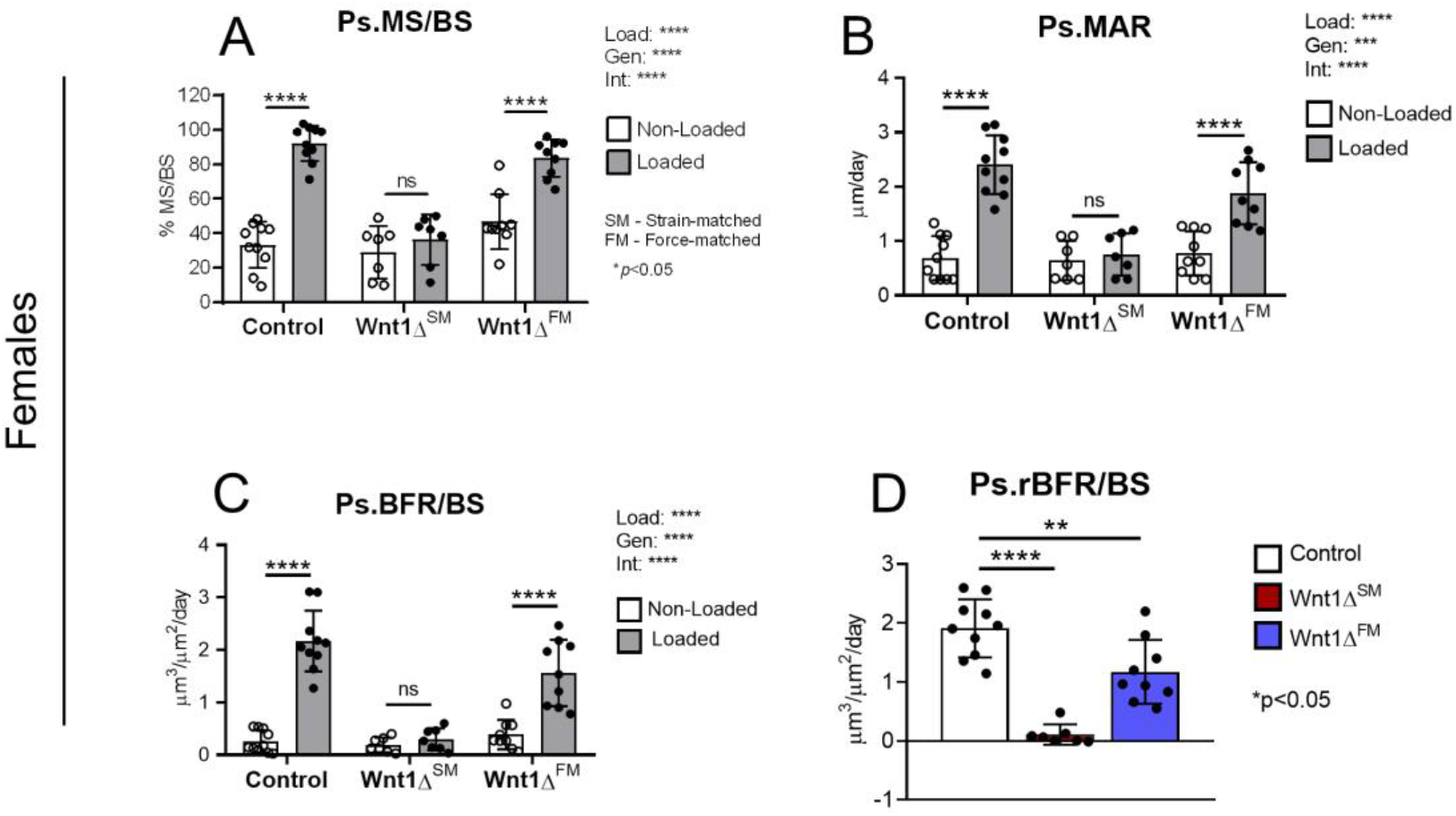
Loading-induced periosteal bone formation was blunted in *Wnt1* knockout females. (A-C) Periosteal bone formation indices in females. Wnt1F/F females were loaded to −9.2N to engender a peak target strain of −3040µƐ. *Wnt1* knockouts were subjected to strain-matched loading (SM) or force-matched loading (FM) by applying either −5.4N or −9.2N, respectively.

To achieve a strain-matched stimulus of −3000µƐ, the force applied to female *Wnt1* knockout mice was 40% less than the force applied to control mice (−5.4N vs. −9.2N; Fig S2), which is consistent with difference in bone morphology (lower Ct.Ar and pMOI in knockouts; Fig 3B, F). Because this level of stimulus did not induce a significant loading response in female knockouts (Wnt1Δ^SM^ group), we asked if a higher magnitude stimulus might induce a response. Thus, we subjected a second cohort of knockout females (Wnt1Δ^FM^) to −9.2N to achieve a “force match” (compared to control); this engendered an estimated −5600µƐ peak strain (Fig S2), a level that typically induces a woven bone response^12^. Knockout females subjected to force-matched loading did display a significant periosteal loading response. Ps.MS/BS, Ps.MAR, and Ps.BFR/BS were 1.8, 2.4 and 4-fold higher, respectively, in loaded vs non-loaded limbs (Fig 5A-5C). Notably, these values were nominally less than the respective fold-increases in control mice (2.8, 3.5 and 8.3), and Ps.MAR and Ps.BFR/BS were significantly less in loaded limbs of force-matched knockout females compared to controls. Moreover, periosteal rBFR/BS was 56% lower in knockouts relative to controls after force-matched loading (*p*<0.01; Fig 5D). Therefore, despite being loaded to 85% higher peak strain the control mice, *Wnt1* knockout females still had a blunted periosteal response to loading.

Finally, in contrast to the robust loading response elicited on the periosteal surface of control mice, tibial loading did not induce a strong endocortical response, which is consistent with the lower values of strain engendered on this surface. In males, indices of endocortical bone formation were not significantly greater in loaded compared to non-loaded limbs, for both control and knockout mice (Fig S4). In female control mice, loading did stimulate a modest increase in Ec.MAR (*p*<0.05, loaded vs. non-loaded) and Ec.BFR/BS (*p*=0.052), but not in strain-matched *Wnt1* knockout mice (Fig S5). Force-matched loading did induce a significant increase in Ec.MAR and Ec.BFR in knockout females, with values in loaded limbs not different from the loaded limbs of control mice.

### Induction of osteogenic genes was diminished in the bones of Wnt1 knockouts

Gene expression analysis was performed 4-hr after the 5^th^ bout of loading. Congruent with previous reports ^11–12^, we found that loading potently induced *Wnt1* expression in the bone (Fig 6A). RT-qPCR analysis indicated that *Wnt1* expression was 6.6-fold higher in the loaded vs non-loaded tibias of Wnt1F/F controls (*p*<0.0001). In contrast, loading had no effect on the abundance of *Wnt1* mRNA in the tibias of *Wnt1* knockouts (*p*=0.999) (Fig 6A), consistent with bone-targeted knockout.

**Figure 6.**
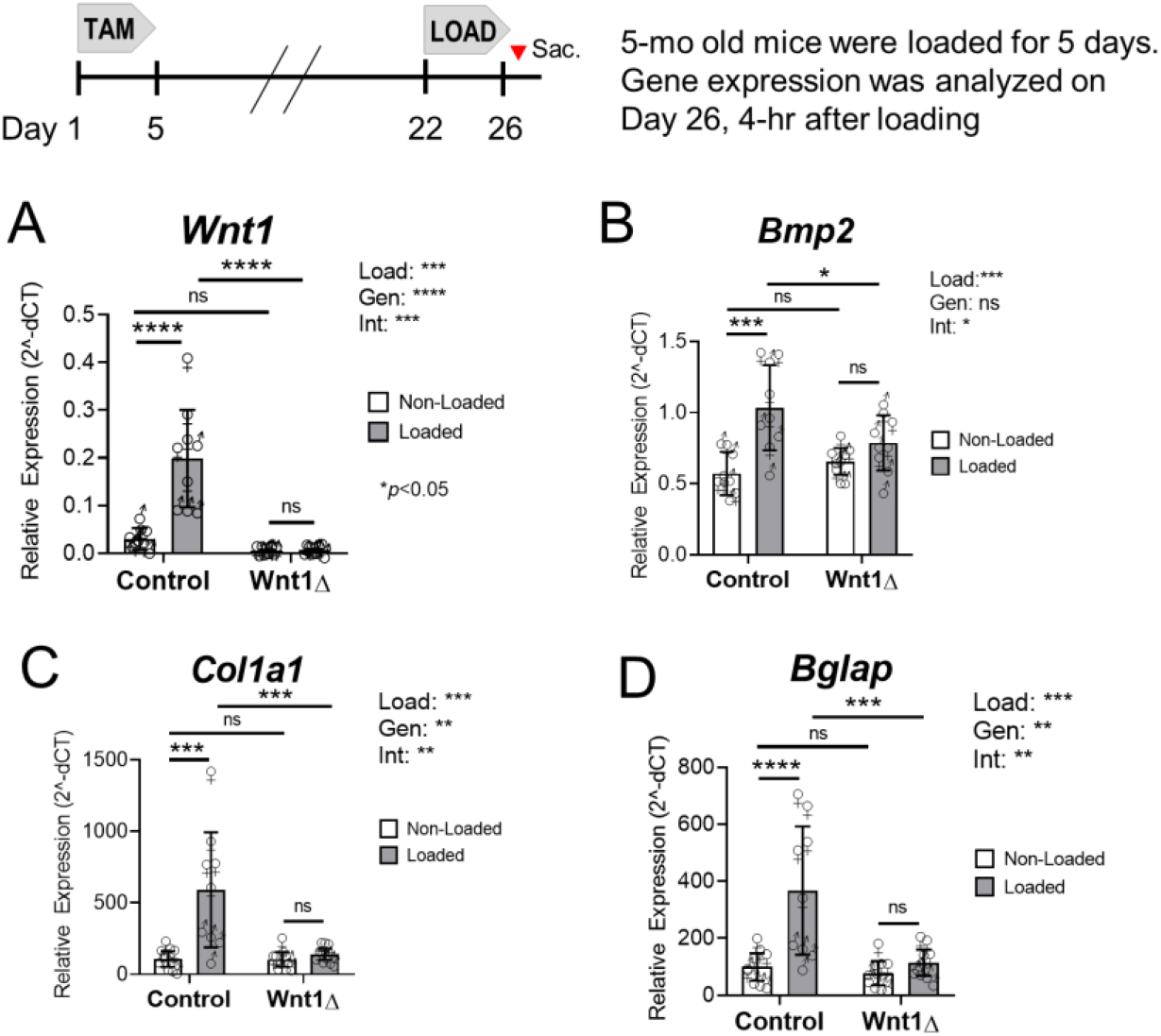
Loading-induced upregulation of bone formation genes was impaired in *Wnt1* knockouts. (A) *Wnt1* induction was observed in Wnt1F/F controls, but not in *Wnt1* knockouts. (B-D) Loading-induced *Bmp2*, *Col1a1*, and *Bglap* upregulation was blunted in *Wnt1* knockouts relative to controls. n=4-5 per sex per genotype in each group for a total of n=9/genotype. Two-factor ANOVA was used to evaluate the effect of tibial loading (“Load)” and genotype (“Gen”), and their interaction (“Int”), on gene expression.

Expression of genes associated with osteo-induction, osteoblast maturation, and matrix synthesis were also analyzed. RNAScope *in situ* hybridization showed that loading increased *Bmp2* expression in osteocytes and *Bglap* expression on the bone surface (Fig S6). These findings were supported by qPCR, which showed that *Bmp2* and *Bglap* were 1.8 and 3.7-fold higher, respectively, in the loaded vs non-loaded bones of Wnt1F/F controls (*p*<0.001) (Fig 6B, 6D). In contrast, loading did not significantly increase *Bmp2* or *Bglap* expression in knockouts. Similarly, loading increased expression of the matrix gene *Col1a1* in the tibias of control mice (5.5-fold higher in loaded vs non-loaded, *p*<0.001), while in knockouts *Col1a1* did not increase with loading (Fig 6C).

Finally, we assayed several reported Wnt-target genes. Neither loading nor genotype had a significant effect on the expression of *Axin2* or *Nkd2* in the bone (Fig S7). In contrast, gene expression analysis by two-factor ANOVA revealed that loading had a modest but significant effect on the expression of *Opg*. In controls, *Opg* was 1.4-fold higher in loaded vs non-loaded limbs (*p*=0.077), while in knockouts *Opg* was only 1.2-fold higher (*p*=0.482) (Fig S7). In sum, these results indicate that *Wnt1* induction in Osx-lineage cells is required for the bone anabolic effects of tibial loading.

### The skeletal phenotype caused by Wnt1 deletion was recapitulated in Wnt1/7b knockout mice

Because *Wnt7b* is osteo-anabolic^14^ and is induced in the bone by loading^12^, we asked whether the phenotypes observed in *Wnt1* knockouts would be exacerbated by also targeting *Wnt7b*. We confirmed recombination of *Wnt1* and *Wnt7b* genes in tibias of *Wnt1/7b* knockout mice (tamoxifen-treated OsxCreERT2; Wnt1F/F; Wnt7bF/F) (Fig.S8A). We also confirmed knockdown of *Wnt1* (−63% in the loaded tibias of Wnt1/7bΔ mice vs control, *p*=0.083) and *Wnt7b* (−87% in the loaded tibias of *Wnt1/7b* knockouts vs control, *p*=0.194) at the mRNA level (Fig S9A).

The cortical bone phenotype observed in *Wnt1* knockouts was recapitulated in *Wnt1/7b* knockouts. In brief, relative to Wnt1F/F; Wnt7bF/F controls, the bone morphology of *Wnt1/7b* knockouts was not different on Day 0 (prior to tamoxifen induction), but on Day 22 cortical bone area, cortical thickness, and pMOI were less in *Wnt1/7b* knockouts. In general, we observed similar magnitude differences in *Wnt1/7b* knockout mice compared to its control, as observed in *Wnt1* knockouts compared to its control. For example, cortical area was 10-15% less in *Wnt1* knockouts vs. control (Fig. 3B), and 15-18% less in *Wnt1/7b* knockouts vs. control (Fig. S8C).

Finally, when *Wnt1/7b* knockout mice and their controls were subjected to strain-matched tibial loading (approx. −3000 με), *Wnt1/7b* knockouts had impaired periosteal bone formation, similar to *Wnt1* single knockouts. In controls, loading significantly enhanced Ps.MS/BS and Ps.MAR, resulting in an overall increase in Ps.BFR/BS of 11 and 5.6-fold in the loaded bones of male and female mice, respectively, compared to non-loaded controls (*p*<0.0001; Figure S9B).

By comparison, loading did not significantly increase bone formation indices in *Wnt1/7b* knockouts. Consequently, the relative bone formation rate (rBFR/BS) in *Wnt1/7b* knockout mice was 90% lower in males and 86% lower in females compared to their respective controls. These data further support the finding that *Wnt1* is a crucial mediator of loading-induced bone formation.

## Discussion

Mechanical loading on the skeleton stimulates bone formation. To investigate the role of *Wnt1* in the skeletal response to loading, we tamoxifen-dosed 5-month old OsxCreERT2; Wnt1F/F mice to inactivate *Wnt1* in the osteoblast lineage, then subjected mice to *in vivo* tibial loading 21 days later. Over 3 weeks, bone area decreased 10-15% in *Wnt1* knockouts, concomitant with a 78% reduction in tibial *Wnt1* expression (Figs 2D, 3B). In Wnt1F/F control mice, loading potently increased *Wnt1* expression in the bone and *in situ* hybridization showed that this induction occurred in osteocytes (Figs 1A, 6A). Loading-induced *Wnt1* upregulation was associated with a 8.4 to 9.6-fold increase in periosteal bone formation rate (Ps.BFR/BS) in control mice (Figs 4D, 5C). In knockouts, Osx-targeted *Wnt1* deletion blocked *Wnt1* upregulation (Fig 6A), leading to a 65 to 94% lower periosteal bone formation response (Figs 4E, 5D). In sum, these findings show that *Wnt1* in the Osx-lineage mediates the bone anabolic effects of skeletal loading in adult mice.

### Loading increases Wnt1 expression in osteocytes

Whole-bone transcriptional profiling studies have shown that mechanical loading increases the expression of Wnt family genes – including *Wnt1* and *Wnt7b* – in the bone^9–10^. Subsequently, Harris and Silva used laser capture microdissection to isolate intracortical bone samples from loaded and non-loaded bones of C57Bl/6 mice, and showed that *Wnt1* in these osteocyte-enriched samples was upregulated by loading^11^. Congruent with these results, our gene expression analyses showed that *Wnt1* was 6.6-fold higher in the loaded vs non-loaded bones of Wnt1^F/F^ mice after 5 days of loading (Fig 6A). Additionally, analysis by RNAscope^®^ *in situ* hybridization showed that *Wnt1* induction occurred primarily in osteocytes embedded in the intra-cortical matrix rather than in cells lining the bone surface (i.e., osteoblasts) (Figs 1A and S1). Analysis of *in situ* hybridization further revealed that loading increased not only the total percentage of *Wnt1*-positive cells in the bone, but also the magnitude of *Wnt1* expression within those cells (Fig 1B). Together, these data show that mechanical loading stimulates *Wnt1* expression in the bone, particularly in osteocytes.

### Loading-induced bone formation is reduced in Wnt1 knockouts

OsxCreERT2 was used to inactivate *Wnt1* in the Osx-lineage cells of 5-month old mice. Previously, we used a similar tamoxifen dosing strategy for Cre reporter analysis in 5-month old OsxCreERT2; Rosa-Ai9 mice, which showed robust TdTomato reporter expression on the bone surface (osteoblasts) and within the cortical bone (osteocytes) within days of tamoxifen treatment, indicating that OsxCreERT2 targets both osteoblasts and osteocytes in the bones of adult mice^7^. OsxCreERT2-mediated *Wnt1* deletion completely neutralized the effect of loading on *Wnt1* expression in the bone (Fig 6A). While tibial *Wnt1* expression increased 6.6-fold in loaded bones of Wnt1F/F controls, *Wnt1* expression remained unchanged in the loaded vs non-loaded limbs of *Wnt1* knockouts. Additionally, tibial loading had a robust inductive effect on the expression of *Col1a1*, *Bmp2*, and *Bglap* in the bones of control but not knockout mice, indicating that loading-induced *Wnt1* expression was required for the regulation of these genes (Figs 6B-C). In combination with our *in situ* hybridization findings, these results suggest that loading stimulates osteoblast differentiation/activity on the bone surface (*Bglap*, Fig S6C) by inducing *Wnt1* ligand expression in osteocytes (Figs 1A, S6A-B).

At the tissue level, *Wnt1* inactivation in the Osx-lineage reduced the periosteal response to loading. In males, periosteal bone formation rate was higher in the loaded vs non-loaded bones of all mice, independent of genotype. However, the relative bone formation rate – which reflects the overall magnitude of the loading response – showed that the periosteal response to loading was 65% lower in knockouts (Fig 4E). In females, strain-matched loading increased all indices of periosteal bone formation in control but not knockout mice (Fig 5A-C). Compared to Wnt1F/F controls, the periosteal response to loading (i.e., Ps.rBFR/BS) was 94% lower in strain-matched *Wnt1* knockouts (Fig 5D). Moreover, analysis of a force-matched cohort (−9.2 N, *Wnt1* knockout females) showed that although a loading response could be elicited with greater strain magnitude (−5600ue vs −3000ue), the periosteal loading response was still 39% less in knockouts vs controls (Fig 5D). Together, these results demonstrate that *Wnt1* is required for loading-induced periosteal bone formation.

### Bone area decreased in Wnt1 knockouts after 21 days deletion

Independent of loading, short-term *Wnt1* deletion (3 weeks) in the Osx-lineage was associated with decreased bone area and cortical thickness (Fig 2B, 2D). These findings are congruent with a report by Wang, et al. which showed that deleting *Wnt1* in the Osx-lineage impaired periosteal bone growth in young mice. Using a Dox-repressible system, Wang et al. showed that inactivating *Wnt1* in the Osx-lineage of 4-week old mice produced measurable deficits in the skeleton 4 and 8 weeks later. Analysis of distal tibias from 12 week old mice revealed that these deficits were caused by a reduction in periosteal bone formation, while changes to endocortical bone formation were not significant^21^.

Similar findings were reported by Joeng, et al. and Luther, et al. Mice lacking *Wnt1* in late osteoblasts (Dmp1Cre; Wnt1F/F) had skeletal deficits – including reduced Ct.Th, Tb.N, and Tb.Th – 2 months after birth^13^, while mice lacking *Wnt1* in osteo-progenitors (Runx2Cre; Wnt1F/F) exhibited BV/TV and Ct.Th deficits in the femur at 24 weeks, leading to increased risk of spontaneous fractures^22^. Hypomorphic *Wnt1* variants have also been liked to low bone mass and spontaneous fractures in mice^23^. Our study extends these findings by showing that short-term *Wnt1* deletion in adults leads to bone loss. Thus, osteoblast/osteocyte *Wnt1* is required for adult bone homeostasis.

The skeletal response to bone anabolic loading declines with age^24–26^. For example, tibial loading enhances periosteal bone formation in adult mice of different ages; however, the magnitude of the osteogenic response to loading is significantly lower in aged 22-month old vs. young adult 5-month old mice^12^. To understand the basis for these differences, Chermside-Scabbo, et al. used RNASeq to characterize the transcriptional responses to loading in 5 and 22-month old C57Bl/6 mice after 1, 3, and 5 days of loading^10^. In both age groups, *Wnt1* emerged as a top 10 upregulated gene on days 1 and 3. The fold-change difference for *Wnt1* in 5-month old mice was 5.8 and 6.7 on days 1 and 3, respectively, while in 22-month old mice the fold-change difference for *Wnt1* was 1.4 and 2.3. Thus, loading increased *Wnt1* in both age groups, albeit to a lesser degree in aged mice – concomitant with a diminished osteogenic response to loading. Taken together, these findings suggest that diminished induction of *Wnt1* by loading may contribute to reduced loading-induced bone formation in the bones of aged mice.

The bone anabolic function of *Wnt1* in the Osx-lineage is further supported by our findings in *Wnt1/7b* double conditional knockout mice, which exhibited cortical bone deficits 21 days post-deletion (Fig S8C). Moreover, like *Wnt1* knockouts, *Wnt1/7b* knockouts did not exhibit a loading-induced increase in *Wnt1* expression and had a blunted periosteal response to loading (Fig S9B). In general, the magnitude of the deficits in *Wnt1/7b* knockouts were comparable to those in *Wnt1* knockouts. Importantly, while we did see evidence of *Wnt7b* gene recombination in *Wnt1/7b* double knockout mice (Fig. S8A), we did not confirm significant knockdown of *Wnt7b* at the mRNA level (Fig. S9A). Thus, we are not able to infer whether *Wnt7b* has a functional role in loading-induced bone formation. Additional studies, perhaps using a different *Wnt7b* floxed mouse, are needed to address this question. Nonetheless, the consistent deficits in loading-induced bone formation in *Wnt1* and *Wnt1/7b* knockout mice show that in two different mouse lines, *Wnt1* in the Osx-lineage was essential for the response to loading.

In conclusion, our findings support a growing body of evidence that *Wnt1* is essential for bone homeostasis in the adult skeleton. In particular, we showed that mechanical loading induced *Wnt1* expression in osteocytes, and that loading-induced *Wnt1* expression in the Osx-lineage was required for the periosteal bone formation response to tibial loading.

## Supplemental Figures

**Figure S1.**
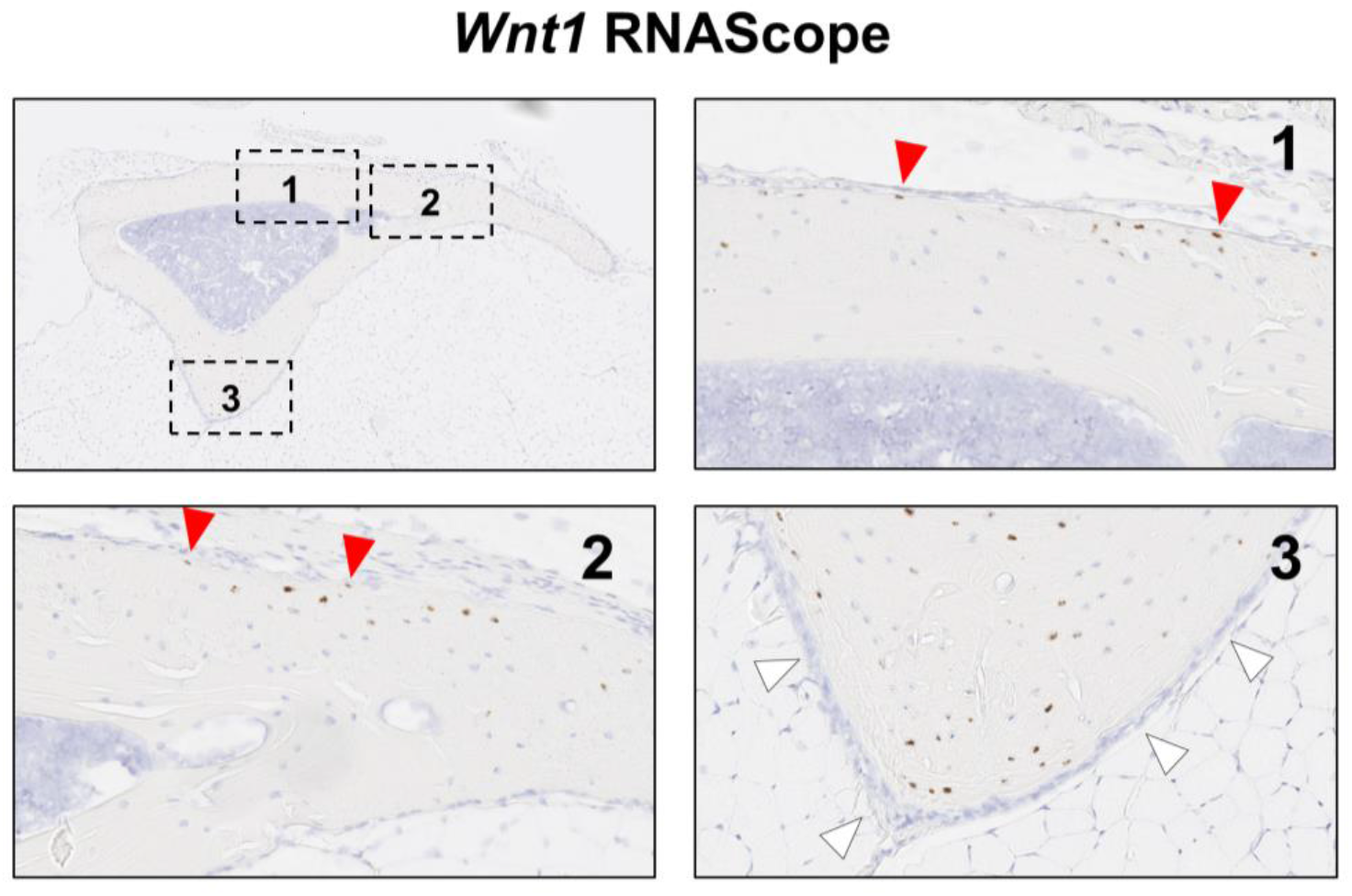
*Wnt1* expression in a loaded tibia after 5 days of loading. *Wnt1* was observed in osteocytes on both the tensile and compressive sides of the bone, including osteocytes near the bone surface (red arrowheads). *Wnt1* expression was not observed in the periosteal layer (white arrowheads). Results representative of n=4.

**Figure S2.**
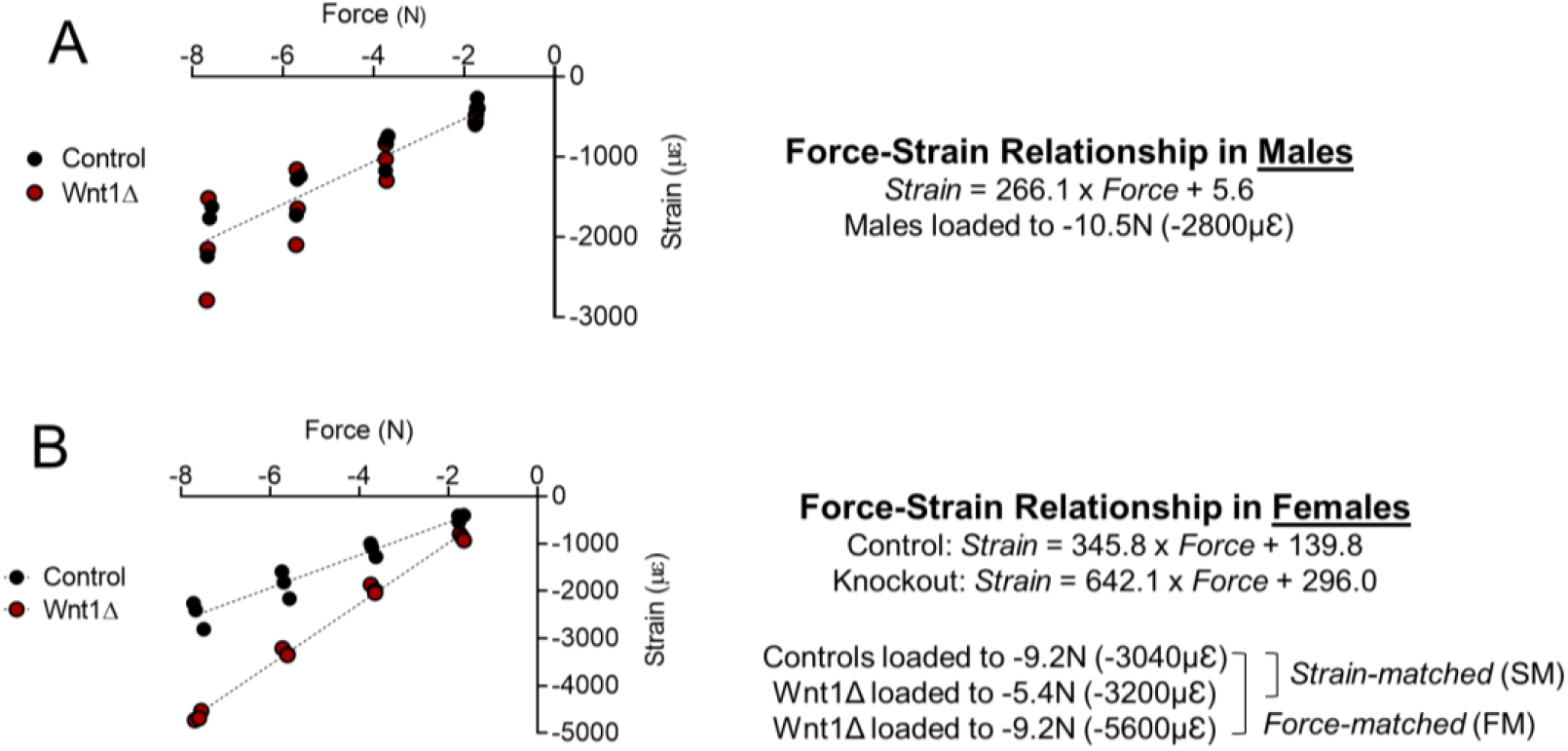
Strain gage analysis was used to experimentally determine the force-strain relationships in male (A) and female (B) mice 3 weeks after tamoxifen induction. In males, strain-matched loading was achieved by loading controls and knockouts to −10.5N, which engendered a peak target strain of −2800µƐ in both genotypes. In females, strain-matched (SM) loading was achieved by loading controls to −9.2N (−3040µƐ) and knockouts to −5.4N (−3200µƐ). Female knockouts in the force-matched (FM) group were subjected to a peak target strain of −5600µƐ.

**Figure S3.**
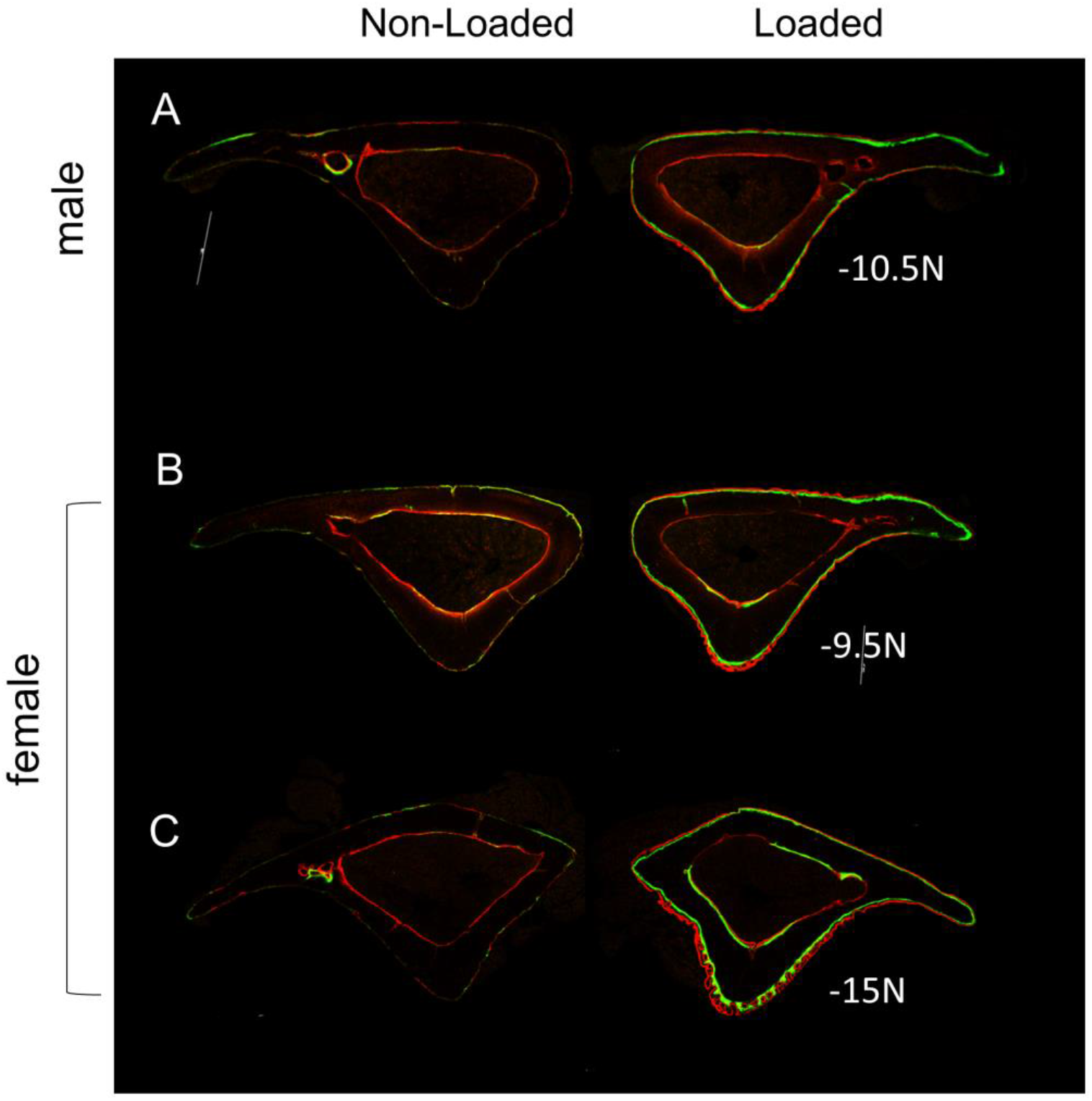
(A-B) Wnt1F/F mice were loaded to a peak target strain of approximately −3000µƐ (−10.5N males; −9.2N females) to stimulate lamellar bone formation. (C) A Wnt1F/F female loaded to −5600µƐ (−15N) had a woven bone formation response to loading. Scale bars represent 500um.

**Figure S4.**
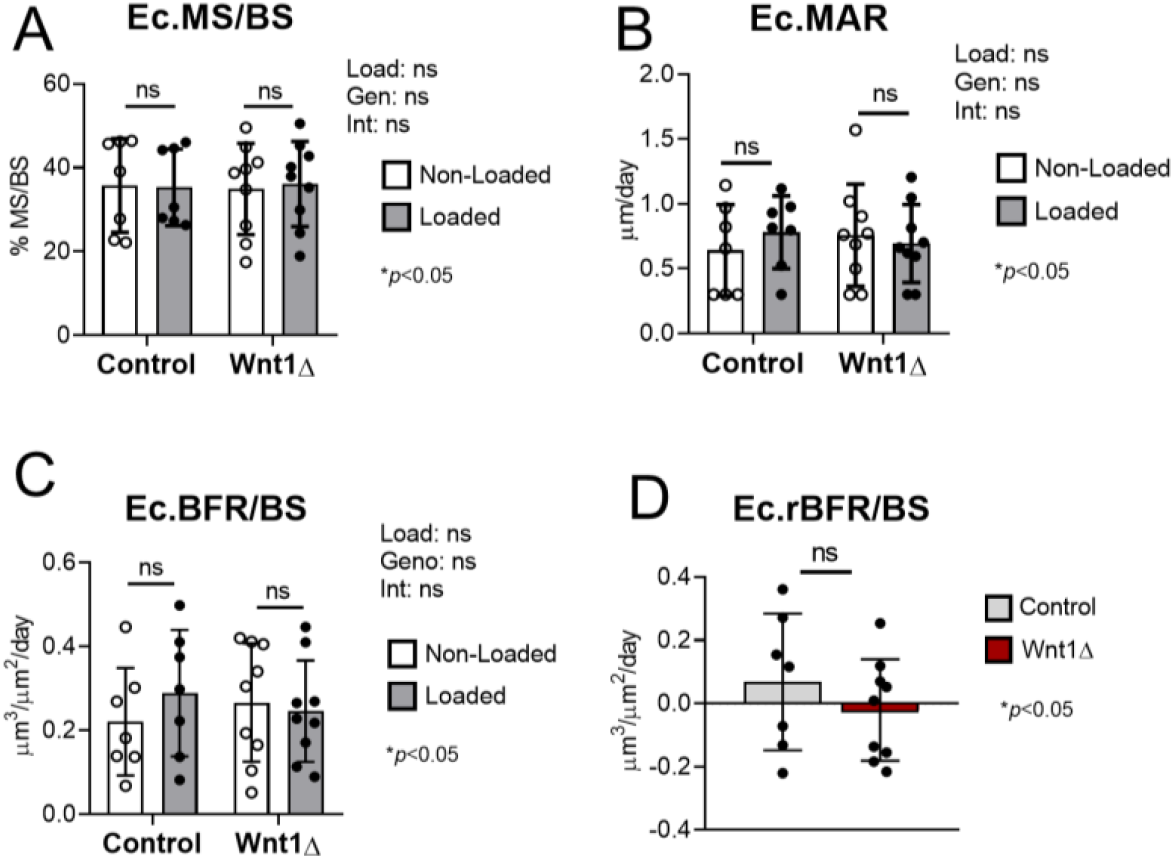
Endocortical bone formation indices in males. In both groups, loading had a negligible effect on endocortical indices.

**Figure S5.**
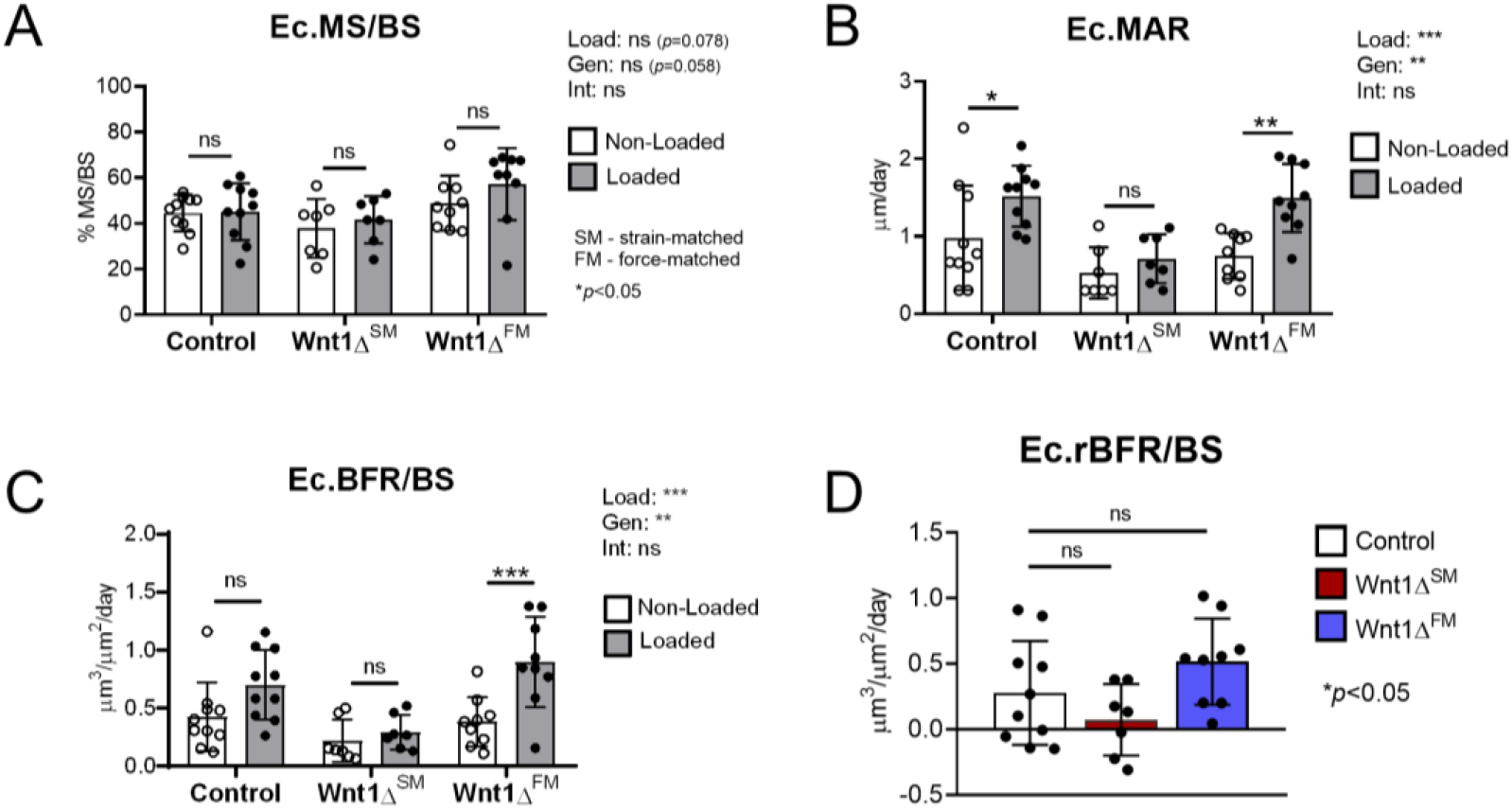
Endocortical bone formation indices in females.

**Figure S6.**
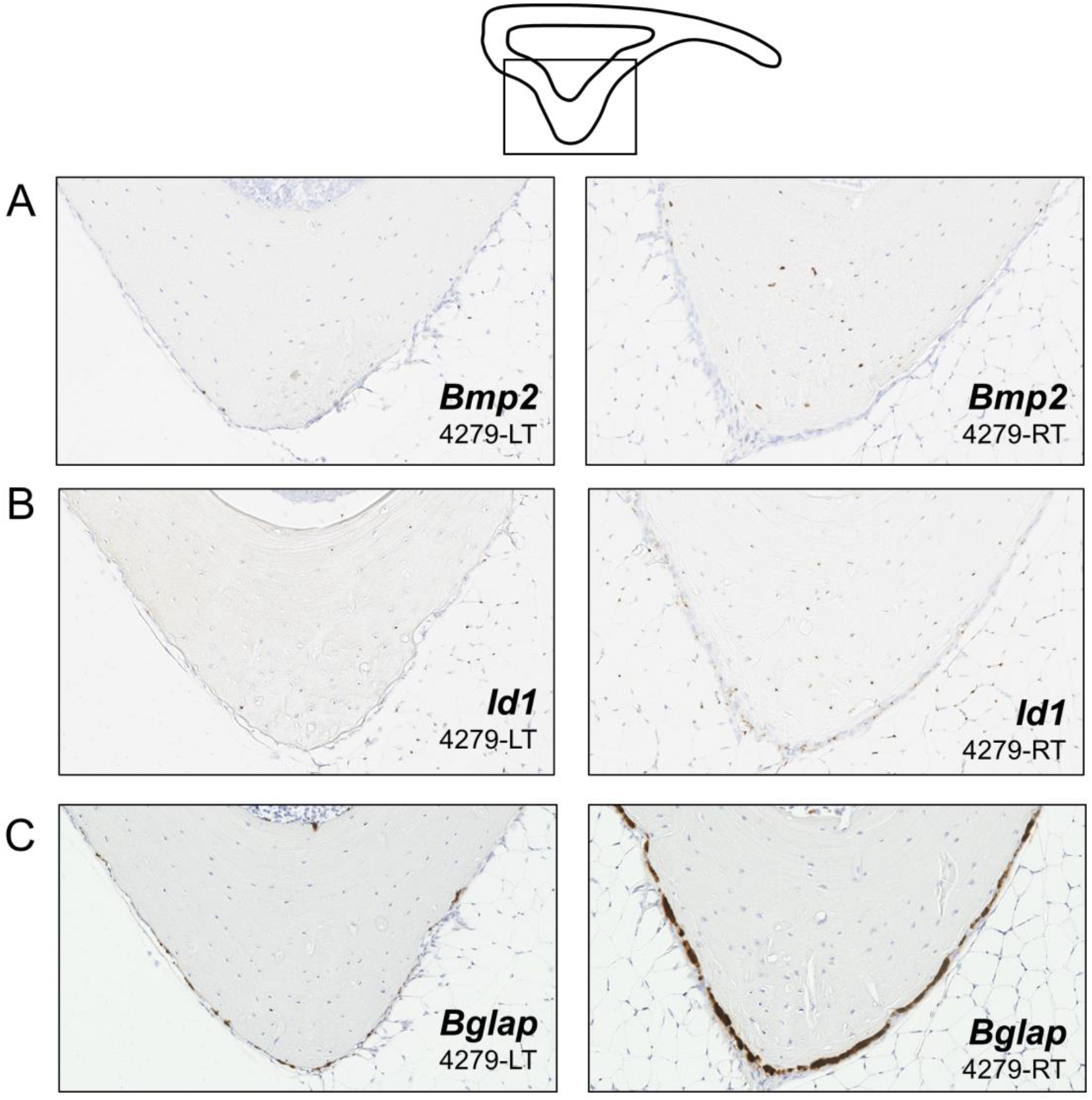
Supplemental RNAScope. After 5 days of loading, bilateral tibias from Wnt1F/F mice were harvested for analysis by RNAScope *in situ* hybridization. (A) Loading increased osteo-inductive *Bmp2* expression in the bone, particularly in osteocytes located at the site of peak compressive strain. (B) Loading also increased the expression of Bmp target gene *Id1* on the bone surface. (C) Robust *Bglap* expression was observed on the surface of loaded bones. Images representative of n=3.

**Figure S7.**
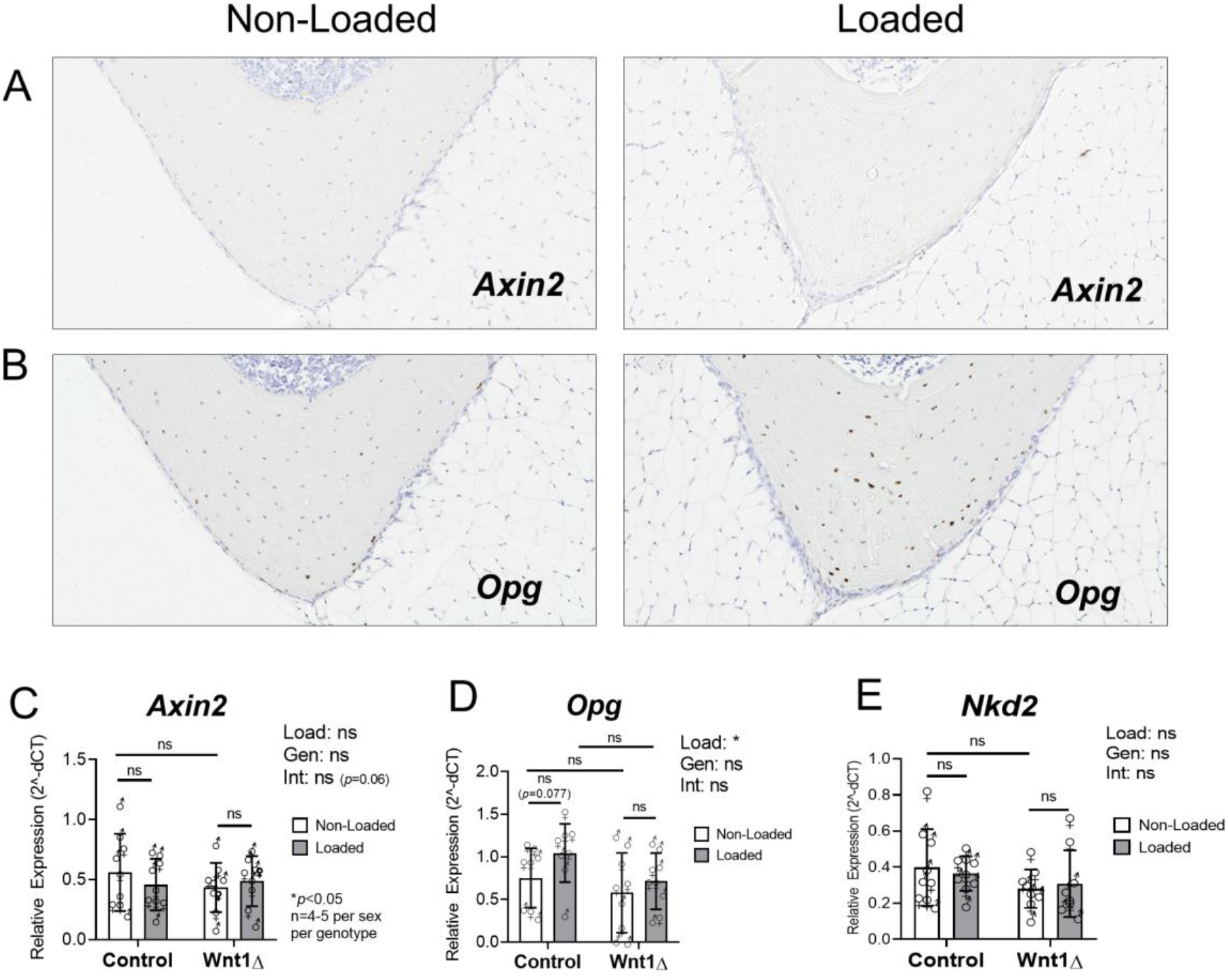
Wnt target genes. Gene expression analysis by RNAScope (A-B) and RT-qPCR (C-E) showed that *Axin2* and *Nkd2* were not different in the loaded vs non-loaded bones of control or knockout mice. Loading increased *Opg* in the tibias of Wnt1F/F controls but not *Wnt1* knockouts. RNAScope images representative of n=3.

**Figure S8.**
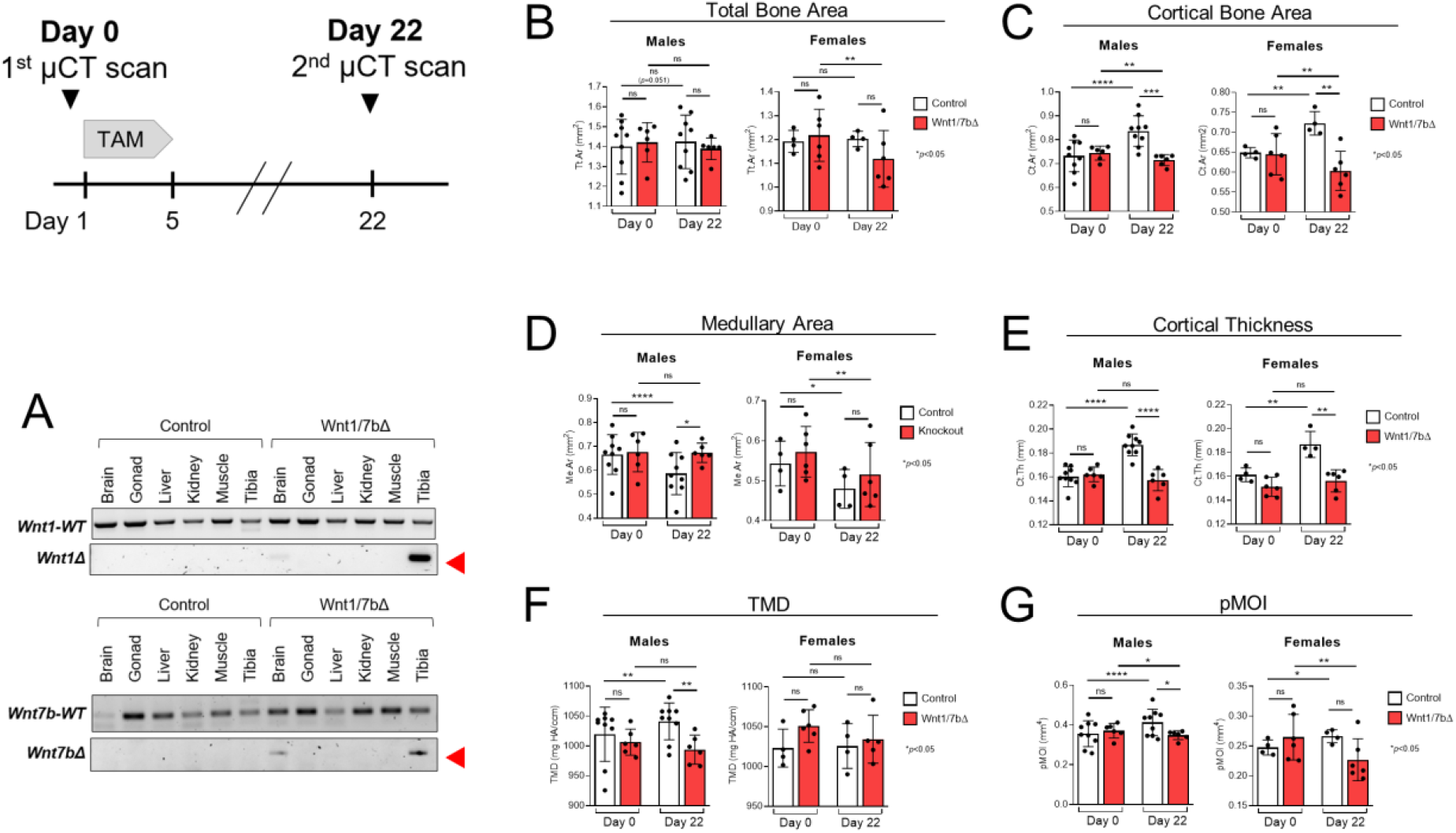
The skeletal deficits associated with *Wnt1* deletion were re-capitulated in *Wnt1/7b* knockouts. OsxCreERT2; Wnt1F/F; Wnt7bF/F (*Wnt1/7b* knockout) mice were tamoxifen-treated at 5-months old to induce *Wnt1* and *Wnt7b* deletion in the Osx-lineage. (A) DNA recombination PCR was performed on Day 22 to confirm bone-specific deletion of *Wnt1* (exon 2-3) and *Wnt7b* (exon 1). (B-G) Serial microCT scans were used to evaluate the effect of *Wnt1*/*7b* deletion (Wnt1/7bΔ) over a 3 week period. Tamoxifen-treated cre-negative Wnt1F/F; Wnt7bF/F mice served as control. n=6-9/group.

**Figure S9.**
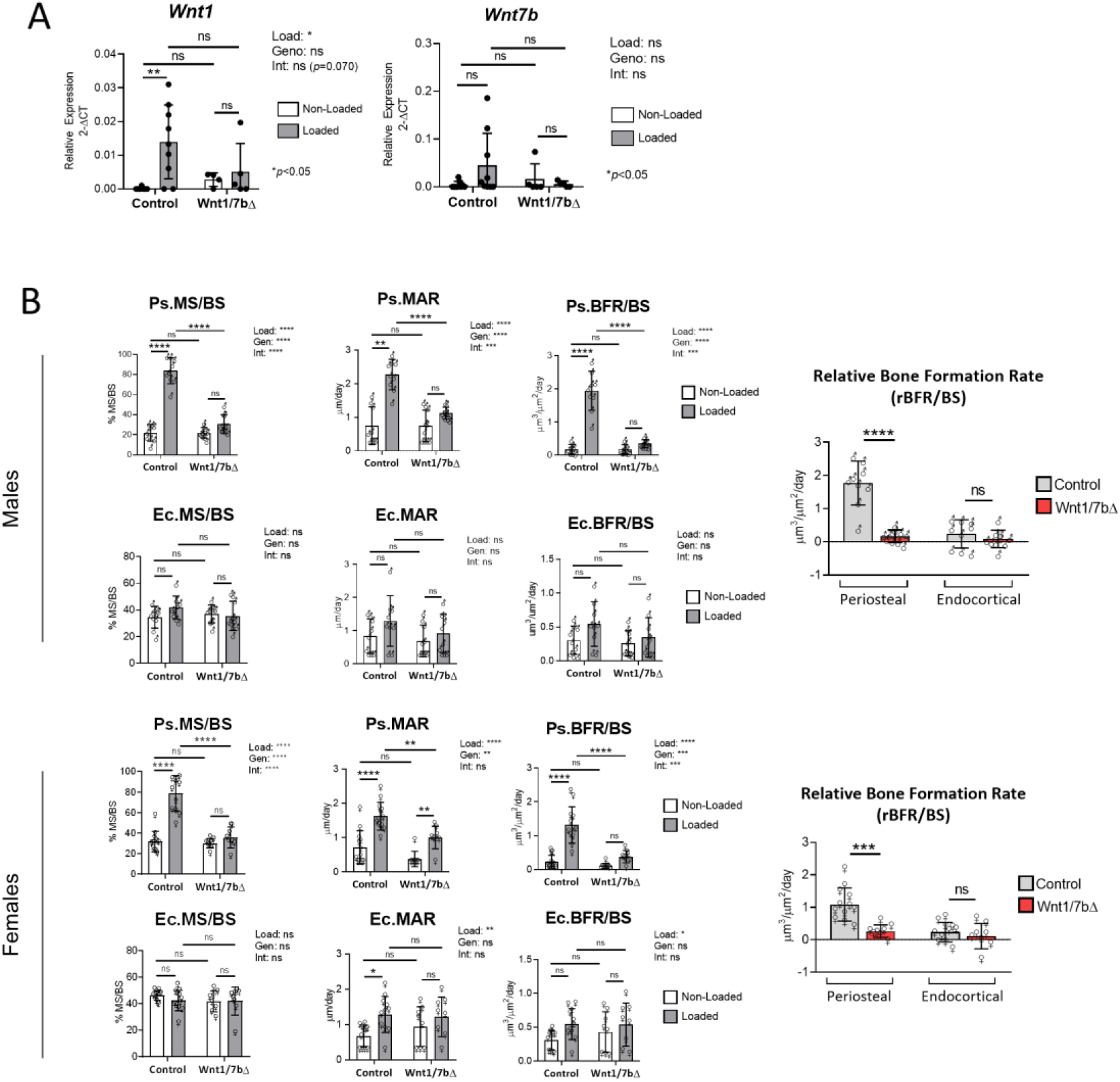
Loading-induced bone formation was reduced in *Wnt1/7b* knockouts. (A) *Wnt1* and *Wnt7b* expression was assayed 4 hours after the 5^th^ bout of loading. (B) Loading-induced periosteal bone formation was reduced in Wnt1/7b knockout male (top) and female (bottom) mice, relative to sex-matched controls.

## Data Availability Statement

The data that support the findings of this study are openly available in Biorxiv at https://doi.org/10.1101/2022.02.28.482178.

## Conflict of Interest Statement

LYL, NM, CCS, JTS, KJ, and BL have no financial or non-financial competing interests to disclose. MJS is on the editorial board at *Bone*, *Journal of Orthopaedic Research*, and *Calcified Tissue International*, and served on the board of directors at the Orthopaedic Research Society (2018-2022). RC serves as Editor-in-Chief for *Journal of Bone and Mineral Research* and has received research support from Ultragenyx.

## Author Contributions

Authors’ roles: Study design: LYL, NM, CCS, and MJS. Study conduct and data collection: LYL, NM, CCS, and JTS. Data analysis and interpretation: LYL, NM, CCS, KJ, RC, BL, and MJS. Drafting manuscript: LYL and MJS.

## Acknowledgments

This work was supported by NIH grants R01 AR047867, T32 AR060719 and the Washington University Musculoskeletal Research Center (P30 AR074992). We thank Crystal Idleburg and Samantha Coleman for histology support.

## Notes

### Competing Interest Statement

The authors have declared no competing interest.

